# Widespread occurrence of benzimidazole resistance single nucleotide polymorphisms in the canine hookworm, *Ancylostoma caninum*, in Australia

**DOI:** 10.1101/2024.09.05.611542

**Authors:** Swaid Abdullah, Thomas Stocker, Hyungsuk Kang, Ian Scott, Douglas Hayward, Susan Jaensch, Michael P. Ward, Malcolm K. Jones, Andrew C. Kotze, Jan Šlapeta

## Abstract

Canine hookworm (*Ancylostoma caninum*), a gastrointestinal nematode of domestic dogs, principally infects the small intestine of dogs and has the potential to cause zoonotic disease. In greyhounds and pet dogs in the USA, *A. caninum* has been shown to be resistant to multiple anthelmintics. We conducted a molecular survey of benzimidazole resistance in *A. caninum* from dogs at veterinary diagnostic centers in Australia and New Zealand. First, we implemented an internal transcribed spacer (ITS)-2 rDNA deep amplicon metabarcoding sequencing approach to ascertain the species of hookworms infecting dogs in the region. Then, we evaluated the frequency of the canonical F167Y and Q134H isotype-1 β-tubulin mutations, which confer benzimidazole resistance, using the same sequencing approach. The most detected hookworm species in diagnostic samples was *A. caninum* (90%; 83/92); the related Northern hookworm (*Uncinaria stenocephala*) was identified in 11% (10/92) of the diagnostic samples. There was a single sample with coinfection by *A. caninum* and *U. stenocephala*. Both isotype-1 β-tubulin mutations were present in *A. caninum*, 49% and 67% for Q134H and F167Y, respectively. Mutation F167Y in the isotype-1 β-tubulin mutation was recorded in *U. stenocephala* for the first known time. Canonical benzimidazole resistance codons 198 and 200 mutations were absent. Egg hatch assays performed on a subset of the *A. caninum* samples showed significant correlation between 50% inhibitory concentration (IC_50_) to thiabendazole and F167Y, with an increased IC_50_ for samples with >75% F167Y mutation. We detected 14% of dogs with >75% F167Y mutation in *A. caninum*. Given that these samples were collected from dogs across various regions of Australia, the present study suggests that benzimidazole resistance in *A. caninum* is widespread. Therefore, to mitigate the risk of resistance selection and further spread, adoption of a risk assessment-based approach to limit unnecessary anthelmintic use should be a key consideration for future parasite control.

## 1. Introduction

For many years the prevailing dogma regarding the use of anthelmintics in small animals such as dogs and cats, as well as in humans, had been that the development of anthelmintic resistance would be slow to occur, or may never occur, or be limited in spread (Schulz et al., 2018; Marsh and Lakritz, 2023). The theory behind this dogma was that, at the time of deworming, the refugia - the population of susceptible nematodes infecting the host - remains substantial (von Samson- Himmelstjerna et al., 2021). By contrast, in ruminants, routine and indiscriminate mass treatment with anthelminthics has led to widespread drug resistance because the refugia has not been retained (Vercruysse et al., 2012; von Samson-Himmelstjerna et al., 2021). However, despite these differences between ruminants and small animals, a number of reports over recent years have indicated that resistance in canine hookworms is indeed a widespread problem in North America (Jimenez Castro et al., 2019; Kitchen et al., 2019; Venkatesan et al., 2023).

Currently, the effectiveness of the benzimidazole anthelmintic class in ruminants varies, with many cases showing reduced efficacy (Ramunke et al., 2016; Jaeger and Carvalho-Costa, 2017; Ploeger and Everts, 2018). This is due to well-documented single nucleotide polymorphisms (SNPs) that result in nematodes carrying alleles resistant to these anthelmintics (Kwa et al., 1993; von Samson-Himmelstjerna et al., 2007). However, this may not apply universally across all countries, production systems, species of nematodes and species of livestock where benzimidazole remains efficacious (Mohammedsalih et al., 2021). In humans, hookworms are commonly treated using benzimidazoles, specifically albendazole and mebendazole (WHO, 2017). Between 1995 and 2015, the efficacy of both albendazole and mebendazole for the treatment of human hookworm infection has decreased by as much as 15%, but resistance as such has not been documented (Schulz et al., 2018).

In dogs, anthelminthic resistance was first documented 20 years ago in hookworms to the pyrimidine derivative anthelmintic pyrantel in Queensland, Australia (Kopp et al., 2007), and currently this anthelmintic is considered ineffective for treating hookworms in dogs in Australia (Dale et al., 2024). However, resistance to benzimidazoles and other anthelmintic classes for dogs in Australia has not been reported. The first report of resistance to benzimidazoles in canine hookworms was from the USA in 2019 (Kitchen et al., 2019). Molecular, phenotypic, and experimental infection evidence demonstrating benzimidazole resistance in *A. caninum* was published independently by two groups in 2019 (Jimenez Castro et al., 2019; Kitchen et al., 2019). These reports first emerged in North America and were followed by an extensive surveillance for the canonical F167Y (T<u>T</u>C>T<u>A</u>C) isotype-1 β-tubulin mutation conferring benzimidazole resistance (Leutenegger et al., 2023, 2024; Venkatesan et al., 2023). A new mutation in isotype-1 β-tubulin at Q134H (CA<u>A</u>>CA<u>T</u>) was experimentally established as another mutation responsible for resistance to benzimidazoles in *A. caninum* and shown to be widespread across North America (Venkatesan et al., 2023).

It is speculated that benzimidazole resistance in *A. caninum* emerged in racing greyhound dogs due to indiscriminate and frequent deworming in racing kennels (Jimenez Castro et al., 2019; Kitchen et al., 2019; Leutenegger et al., 2024). Evidence of multiple anthelmintic drug resistance (MADR) in *A. caninum* led the American Association of Veterinary Parasitologists to establish a hookworm task force to address this issue in 2021 (von Samson-Himmelstjerna et al., 2021; Marsh and Lakritz, 2023). The screening of thousands of dogs positive for hookworms in the USA and Canada demonstrated that the canonical F167Y (T<u>T</u>C>T<u>A</u>C) isotype-1 β-tubulin mutation was present in 11.3% (715/6,329) of faecal *A. caninum*-positive samples (Leutenegger et al., 2024). In a smaller study of samples from the USA, 49.7% (156/314) of the individual samples tested positive for the F167Y (T<u>T</u>C>T<u>A</u>C) mutation (Venkatesan et al., 2023). Reanalysis of next-generation sequence data from Venkatesan et al. (2023) for the presence of non-*A. caninum* sequences revealed the presence of other hookworms (*Ancylostoma braziliense*, *Ancylostoma tubaeforme* and *Uncinaria stenocephala*), but only the benzimidazole-susceptible residues of isotype-1β-tubulin were found in these species (Stocker et al., 2024).

The aim of this study was to investigate the extent of benzimidazole resistance SNPs in canine hookworm (*A. caninum*) outside North America, specifically in Australia and New Zealand. We utilised opportunistic sampling from four different sources in Australia and New Zealand. All hookworm-positive samples were screened using internal transcribed spacer (ITS)-2 rDNA PCR and deep amplicon sequencing to determine the hookworm species present. To investigate resistance to benzimidazoles, we used deep amplicon sequencing to evaluate the nucleotide sequence of the isotype-1 β-tubulin mutations that are known to be associated with resistance to this drug class, specifically the regions that encode amino acids: glutamine 134 (Q134), phenylalanine 167 (F167), glutamic acid 198 (E198) and phenylalanine 200 (F200). We then undertook egg hatch assays (EHAs) on a subset of the samples to correlate the molecular signatures, specifically frequency of F167, with the benzimidazole-resistant phenotype of these field isolates.

## 2. Materials and methods

### 2.1. Ethics statement

The use of residual clinical samples was in accordance with the University of Sydney (Australia) Animal Ethics Committee Protocol 2024/2508. Dog faecal samples for larval cultures were obtained in accordance with the University of Queensland (Australia) Animal Ethics Committee Protocol 2021/AE001169 and the faecal egg count reduction test on one dog was conducted in accordance with the University of Queensland Animal Ethics Committee Protocol SVS/111/20.

### 2.2. Faecal samples with hookworm infection

Faecal samples (*n*=64) were provided by registered veterinary practitioners in accordance with the Veterinary Practice and Animal Welfare Acts, Australia for the purpose of diagnostics used in decision making by the veterinary practitioner. Samples were submitted for parasitological diagnostics (*n*=50) to Vetnostics (Laverty Pathology, North Ryde Laboratory, Sydney, New South Wales, Australia) and (*n*=14) Veterinary Pathology Diagnostics Services (VPDS, The University of Sydney, New South Wales, Australia) for centrifugal qualitative flotation using saturated salt (NaCl, specific gravity = 1.21); refrigerated residual faecal samples that previously returned a positive result for hookworms at the clinical diagnostic laboratory were received for molecular species characterisation. These samples came from 63 dogs and one cat. The group of dogs included 45 greyhounds, four foxhounds, one boxer, one Labrador retriever, one Afghan, one German Shepherd, one King Charles Cavalier, one Irish wolfhound cross, and eight dogs of unknown breed. Material used for hookworm species identification and benzimidazole susceptibility, and subsequent results, were de-identified.

Material (*n*=22) from Queensland, Australia was obtained from dogs from adoption and shelter facilities that receive rescued or surrendered animals from greyhound rescue organisations and pounds across south-eastern Queensland. These sample were from 16 greyhounds, one Siberian husky, three cross-breed dogs and two dogs of an unknown breed. Dogs within these facilities are kept in individual pens, which are cleaned and disinfected daily. Fresh faecal samples were collected from each dog after natural defecation and screened for hookworm infection using faecal flotation as described in section 2.3.

Samples (*n*=20) from Massey University, Palmerston North, New Zealand that contained hookworm eggs were processed to isolate the eggs, which were stored in 80% ethanol for transfer to VPDS.

Details of samples used in this study, with available metadata, are available at LabArchives (https://dx.doi.org/10.25833/hmvy-jn79).

### 2.3. Faecal egg counts (FECs), hookworm egg recovery and DNA isolation

Faecal egg counts (FECs) were performed with saturated salt (NaCl, specific gravity = 1.21) solution using a Whitlock Universal McMaster four chamber worm egg counting slide in two different laboratories.

At the University of Sydney, 3 g of faecal samples were homogenised with 60 ml of saturated salt solution and filtered via a coarse strainer. We counted two compartments each of 0.5 ml volume, resulting in minimum detection of 20 egg per gram of faeces (EPG) (absence of eggs indicates <20 EPG in the sample). Only strongyle-like (=hookworm) eggs were counted. The remainder of the filtered homogenate was poured into a 50 ml Falcon tube and centrifuged at 800 *g* for 10 min for centrifugal floatation. The uppermost supernatant (∼10 ml) was collected with a plastic Pasteur pipette into a new clean 50 ml Falcon tube, topped with tap water and centrifuged at 800 *g* for 10 min. The supernatant was decanted and the pellet that contained the concentrated eggs was retained. Then ∼1 ml of tap water was added with a clean plastic Pasteur pipette to disturb the pellet by repeated aspiration until a homogeneous solution formed. The homogenate was transferred to a clean 1.5ml Eppendorf tube and spun at 8,000 *g* for 10 min. The supernatant was completely removed and the Eppendorf tube with the egg pellet was stored at -20 °C until further use.

At the University of Queensland, 3 g of faecal samples were homogenised with 42 ml of saturated salt solution, filtered through a tea strainer and the filtrate was immediately pipetted into the counting chamber (four compartments each of 0.5 ml volume). Counting all compartments (2 ml) delivered minimum detection of seven EPG (absence of eggs indicates <7 EPG in the sample). Eggs were purified for use in the EHA (see section 2.6). The remaining eggs were suspended in distilled water in 1.5 ml microtubes, incubated for 72 h at 28°C and allowed to hatch. The hatched larvae were stored at -20°C until further use.

All total genomic DNA was isolated at the University of Sydney. For partially cleaned hookworm eggs, an Isolate II Fecal DNA Kit (Meridian Bioscience, Australia) was used according to the manufacturer’s instructions with the following homogenisation step. For purified eggs, 750 µL of lysis buffer were added to the Eppendorf tube with the pellet and homogenised by repeated aspiration with a pipette, then all the contents were transferred into a bead-beating tube, and the eggs disrupted and homogenised using a high-speed benchtop homogeniser, FastPrep-24 (MP Biomedicals, Australia), for 40 s at 6.0 m/s. Every 11^th^ sample was a blank sample that only contained the lysis buffer. Egg samples suspended in ethanol were first dried before processing as described above. For hatched larvae, a Monarch Genomic DNA Purification Kit (New England Biolab, Australia) was used according to the manufacturer’s instructions. Purified genomic DNA samples and blanks were isolated into 100 µL of elution buffer (10 mM Tris-Cl, pH = 8.5) and stored at −20 °C until further use.

### 2.4. Amplicon metabarcoding and Illumina deep sequencing

All isolated DNA samples were first screened for the presence of nematode DNA using nematode ITS-2 rDNA quantitative PCR and samples with a cycle threshold (Ct)<35 were considered suitable DNA material for follow-up processes.

Total DNA was subjected to three independent real-time PCR amplifications at the University of Sydney as previously described (Stocker et al., 2023). ITS-2 rDNA (ITS-2) was amplified by the NC-1 and NC-2 primers designed by Gasser et al. (1993), and two isotype-1 β-tubulin regions were amplified using assays BZ167 and BZ200 developed by Jimenez Castro et al. (2019), which include key diagnostic SNPs conferring benzimidazole resistance. All primers were adapted for Illumina metabarcoding using Next Generation Sequencing (NGS). The ITS-2 target was amplified using 32-36 amplification cycles, while 40 amplification cycles were used for isotype-1 β-tubulin (BZ167, BZ200). All reactions utilized SYBR-chemistry using SensiFAST™SYBR® No-ROX mix (Meridian Bioscience, Australia). Each real-time PCR mix (25 μL) included 2 μL of template DNA (neat) and primers at final concentrations of 400 nM. All PCRs were prepared using a Myra robotic liquid handling system (Bio Molecular Systems, Australia). Reactions were run on a CFX96 Real-Time PCR Detection System with corresponding CFX Manager software (BioRad, Australia). The protocol involved an initial denaturation at 95 °C for 3 min, followed by 32 or 40 cycles of 95 °C for 5 s, 60 °C for 15 s, 72 °C for 15 s, and a final melt curve analysis. Each run included a no-template control (ddH_2_O) to monitor for potential contamination. Amplicons in 96 well plate format were submitted for indexing, purification and amplicon sequencing using Illumina NGS at the Ramaciotti Centre for Genomics, University of New South Wales, Sydney, Australia and sequenced on MiSeq using MiSeq Reagent Kits v2 (250PE, Illumina). De-multiplexed FastQ files were generated via BaseSpace (Illumina).

### 2.5. Analysis of the Illumina sequenced amplicons

All FastQ files were processed through R package ‘dada2’ v1.26.0 (Callahan et al., 2016) in R v4.2.2 (R Core Team, 2022), to obtain amplicon sequence variants (ASV) and their abundance per sequenced amplicon. The dada2 pipeline at the University of Sydney includes filtering, error estimation and denoising, before paired-end reads were merged to reconstruct the full ASV before removal of chimeras. Prior to processing through the dada2 pipeline, primers were removed from all forward and reverse FastQ reads using ‘cutadapt’ version 4.0 (Martin, 2011).

The assignment of species based on ITS-2 ASVs was carried out in CLC Main Workbench v22.0 (Qiagen Australia) as previously described (Stocker et al., 2024). All ASVs were imported into CLC Main Workbench and top blastn hits against the ‘nr’ database at NCBI [https://blast.ncbi.nlm.nih.gov/Blast.cgi] were identified. ITS-2 sequences were discarded for any ASVs where the top blastn hit was not hookworm (Ancylostomatidae). For species assignment we constructed alignment of AVSs with an in-house curated reference ITS-2 hookworm sequence library consisting of publicly available sequences (GenBank: **<u>KP844736</u>**; **<u>MT345056</u>**; **<u>JQ812692</u>**; **<u>LC036567</u>**; **<u>JX220891</u>**; **<u>EU344797</u>**; **<u>AB793527</u>**). The resulting multiple sequence alignment was subject to phylogenetic reconstruction, where species identity was based on manual assignment of ASV to a species based on its position on the phylogenetic tree relative to the reference ITS-2 sequences.

For the β-tubulin sequence data (BZ167, BZ200 assay), the assignment of species was performed with the aid of CLC Main Workbench v22.0 (Qiagen Australia) as previously described in Stocker et al. (2024). Briefly, all ASVs were imported into CLC Main Workbench and top blastn hits against the ‘nr’ database at NCBI were identified. All ASVs where the top blastn hit was a hookworm (Ancylostomatidae) β-tubulin were retained for further analysis. The reference hookworm alignments of BZ167 and BZ200 amplicons according to Stocker et al. (2024) were used to unambiguously identify ASV identity and only those ASVs that were confirmed based on phylogenetic analysis were included in further analyses. The sequence alignment included β-tubulin reference sequences from *A. caninum* (**DQ459314**) (Schwenkenbecher et al., 2007), *Ancylostoma duodenale* (**EF392850**), *U. stenocephala* (Stocker et al., 2023), and *Necator americanus* (**EU182348**), which enabled us to identify the exon/intron boundary and residues that code for susceptibility to benzimidazoles, amino acids amino acids Q134, F167, E198 and F200 (Venkatesan et al., 2023). Our reference alignment included sequence data for isotype-1 β-tubulin from *A. caninum*, *Ancylostoma ceylanicum*, *A. duodenale* and *N. americanus* at ‘WormBase ParaSite’ (Howe et al., 2017; Bolt et al., 2018; Harris et al., 2020).

Following the dada2 pipeline, species assignment raw read counts were converted into relative abundance (percentages). Any species read counts with <25 reads were disregarded as spurious. Proportions of SNPs for amino acids Q134, F167, E198 and F200 were calculated per species for each of the amplicons of isotype-1 β-tubulin (BZ167, BZ200).

The number of samples, and number of *A. caninum* samples with Q134H (CA<u>A</u>>CA<u>T</u>) and F167Y (T<u>T</u>C>T<u>A</u>C) separately, were summed per postcode in New South Wales. Proportional symbol maps were created by joining this data to postcodes of Australia shapefile (Geocentric Datum of Australia, GDA 1994) in ArcGIS^®^ Pro. Spearman rank correlations (r_SP_) were calculated between EPG per sample and both frequency of Q134H (CA<u>A</u>>CA<u>T</u>) and F167Y (T<u>T</u>C>T<u>A</u>C). In addition, the EPG between resistant versus not resistant allele status for both mutations was assessed using a Kruskal-Wallis non-parametric analysis of variance. Spearman rank correlations were also calculated between EPG, frequency of Q134H (CA<u>A</u>>CA<u>T</u>), frequency of F167Y (T<u>T</u>C>T<u>A</u>C) and age (years). Finally, the association between breed (greyhound versus other) and EPG, resistance allele frequency for Q134H (CA<u>A</u>>CA<u>T</u>) and F167Y (T<u>T</u>C>T<u>A</u>C), was assessed using a Kruskal-Wallis non-parametric ANOVA; DFn: degrees of freedom for the numerator of the F ratio, DFd: degrees of freedom for the denominator of the F ratio. The type I error for all statistical tests was set at 0.05 (IBM SPSS Statistics v 28).

### 2.6. In vitro EHA

Eggs were harvested from fresh faecal samples from hookworm-positive dogs for use in in vitro EHAs. Saturated sugar solution (specific gravity = 1.30), three times the weight of individual faecal samples, was added to the faecal samples in a shaker bottle. The bottle was manually agitated to form a slurry, which was then filtered through a 200 µm sieve into a beaker to remove large debris. All of the filtered solution was then transferred into 50 ml Falcon tubes, filling each tube up to 40 ml, and then 10 ml of distilled water were slowly and carefully poured along the edge of the tube to avoid mixing with the faecal solution. Each tube was then screw capped and centrifuged at 400 *g* for 10 min. Following centrifugation, the ‘fuzzy’ layer (water-sucrose interface) of each tube was removed with a pipette and pooled into a 200 mL beaker. The solution was passed through a 75 µm sieve, retaining the filtrate, which was further passed through a 20 µm sieve, which retained the hookworm eggs. This sieve was then retrograde washed with 20 mL of distilled water into a glass Petri dish. The recovered eggs were carefully poured into a two 15 mL centrifuge tubes, leaving the debris in the Petri dish. The eggs were concentrated into 2 mL of water by centrifugation for 5 min at 400 *g* and the supernatant discarded. The egg concentration was estimated by thoroughly mixing the 2 mL of egg solution and counting the eggs in three 20 μl aliquots. The egg numbers were adjusted to approximately two eggs per microlitre either by dilution with distilled water or by concentration using centrifugation as described above.

The EHA was performed using 96-well plates containing thiabendazole embedded in agar, as described previously for assays with human hookworms (Kotze et al., 2009). A stock solution of thiabendazole (5 mg/mL, Sigma-Aldrich Australia) was prepared in dimethyl sulfoxide (DMSO) and serially diluted two-fold in the same solvent. Aliquots (2 μL) from a series of dilutions were added to 96-well microtiter plates, such that each row of the plate comprised a double dilution, ranging from 0.003 μg/mL up to 1.359 μg/mL or 0.011 μg/mL to 5.437 μg/mL. The first two wells of each row were used as control wells (received 2 μL of DMSO only). Molten agar (200 μL at 2% w/v) (Davis Gelatine, Australia; powdered agar Grade J) was dispensed into each well of the plate and allowed to set. Plates were placed into plastic press-seal bags stored at 4°C.

Prior to establishing the assays, the plates were allowed to warm in an incubator set to 28°C. Thirty microlitres of concentrated hookworm egg solution (∼2 eggs/μL) were pipetted into each well. Plates were incubated for 48 h at 28°C, whereupon the assay was terminated by addition of Lugol’s iodine solution. The numbers of unhatched eggs and larvae in each well were counted with the aid of a stereo microscope (Stemi 508, Carl Zeiss, Australia). All assays were performed in triplicate for the sample isolated from each dog sample.

The number of larvae in each well was expressed as a percentage of the total numbers of larvae and unhatched eggs to give the hatch rate in each well, and then this was expressed relative to the mean hatch rate in the control wells. The data were then analysed by non-linear regression using GraphPad Prism 10.3.0 software (GraphPad Software Inc. USA). This analysis yielded 50% inhibitory concentration (IC_50_) values representing the concentration of thiabendazole that resulted in an egg hatch of 50% compared with control assays. The IC_50_ values for the dogs examined in this study were compared with those reported previously for drug-susceptible *A. caninum* isolates: 0.046 and 0.048 µg/mL (Jimenez Castro et al., 2019) and 0.045 µg/mL (Diawara et al., 2013). An Australian fenbendazole-susceptible isolate of *A. caninum* (‘Dougie’, from February 2021) *s*erved as a local positive control and demonstrated faecal egg reduction of 100% 10 days post administration from an initial 525 EPG.

The left-over eggs were suspended in distilled water in 1.5 ml microtubes, incubated for 72 h at 28°C and allowed to hatch. The hatched larvae were stored at - 20°C and subjected to DNA isolation and follow up processes described earlier (section 2.3).

### 2.7. In vivo efficacy measurements

One dog (sample D6) was treated by the shelter facilities and the FEC monitored throughout treatment. The dog was treated according to its weight and the product label on day 0 with Drontal Allwormer (Vetoquinol, Australia; actives: febantel, praziquantel and pyrantel embonate) and on day 21 with Credelio Plus (Elanco, Australia: milbemycin oxime and lotilaner). In animals, febantel is a prodrug that is actively hydrolyzed to the active metabolite fenbendazole. Egg counts were conducted within the period 10-21 days after Drontal Allwormer treatment to calculate the faecal egg reduction percentage (>95% reduction is considered an effective product). Only a single sample was obtained at day 8 after Credelio Plus because the dog was then rehomed.

### 2.8. Data availability

The EHA raw data, FEC reduction data and metadata for all samples are available at LabArchives (https://dx.doi.org/10.25833/hmvy-jn79). Raw FastQ sequence data was deposited at SRA NCBI BioProject: **<u>PRJNA1149426</u>**.

## 3. Results

### 3.1. Dominance of A. caninum across south-eastern Australia using metabarcoded ITS-2 rDNA deep sequencing in dogs

We used 106 faecal samples that were considered hookworm-positive. There were 20 dog samples from New Zealand. For 61 of the 106 samples the worm FEC ranged from 20 to 4,920 (average 526) EPG. The majority of the Australian samples came from a commercial diagnostic laboratory (*n*=50) and were collected over a period of 13 months between March 2023 and March 2024.

There were 86 samples from south-eastern Australia, 85 samples from dogs and one from a cat (TS42). Amplifiable DNA was obtained from 98% (84/86), yielding a Ct<35 (11.8 to 31.6) at ITS-2 in an rDNA assay targeting nematode DNA. All 84 samples yielded metabarcoded and deep sequenced ITS2 rDNA sequence data for nematode species identification (*n*=84; average 21,323 reads, range 794 to 32,899 reads per sample) (Fig. 1A). Two samples yielded a low (<2,000) number of ITS-2 rDNA sequence reads, 794 (TS100) and 1,537 (TS95); removing those the remaining samples yielded >2,000 reads per sample (*n*=82; average 21,815, range 2,736 to 32,899). The cat hookworm sample contained sequences that matched *Ancylostoma tubaeforme* (TS42). The remaining dog samples (*n*=81) contained sequences of *A. caninum* (*n*=77, 95%) and *U. stenocephala* (*n*=4, 5%). One sample (TS35) had both *A. caninum* (96%) and *U. stenocephala* (4%) ITS-2 rDNA sequences. The additional two samples with <2,000 ITS-2 rDNA sequence reads (TS95, TS100) included only *A. caninum* sequences.

**Fig. 1.**
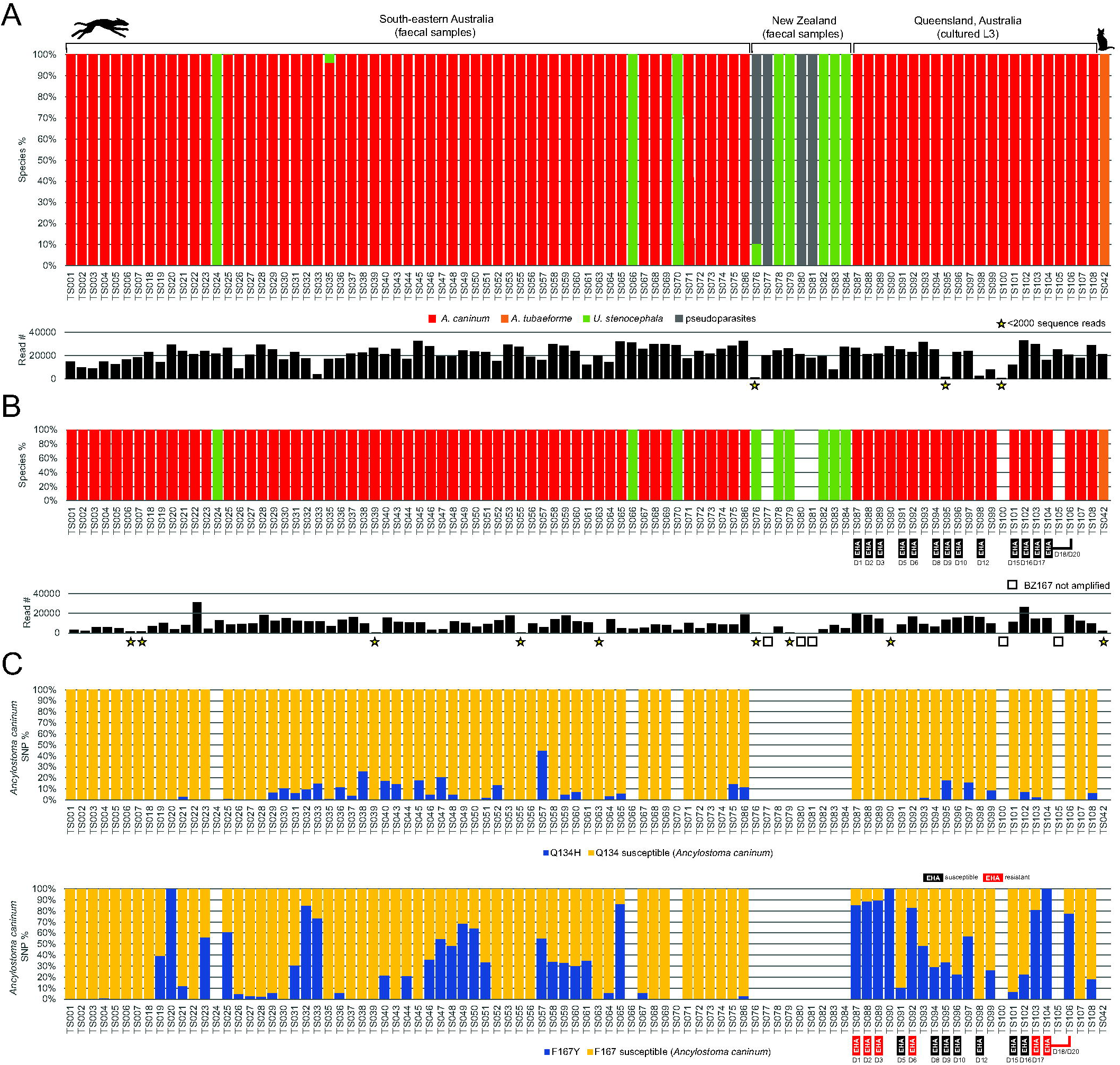
Molecular profiling of hookworm species using internal transcribed spacer (ITS)-2 rDNA and benzimidazole susceptibility using isotype-1 β-tubulin. (A) Percentage of hookworm species within each sample (depicted as percentage) and number of sequencing reads per sample in the bar chart directly below. Sampled with hookworm-like stages that were profiled using metabarcoded deep Illumina amplicon sequencing at ITS-2 rDNA. All but one sample came from dogs. The single, cat sample (TS042) is on the very right. The broad scale of the samples is indicated above the bar chart. Amplicon sequence variants (ASVs) were pooled according to species of nematodes to which they belonged using a reference sequence alignment. Proportion of each species (*Ancylostoma caninum*, *Ancylostoma tubaeforme, Uncinaria stenocephala*) is colour coded in the stacked bar chart; ASVs that belonged to unrelated nematodes are depicted as ‘pseudoparasites’ and represented spurious parasite sequences, specifically trichostrongylid species from sheep and rabbits. The bar chart underneath represents the number of sequencing reads for each sample, and samples with <2,000 sequence reads are depicted with a star. Samples are vertically aligned across the bar charts. (B) Dog samples profiled for hookworm species identification using the BZ167 amplicon of the isotype-1 β-tubulin, specifically the region covering amino acids Q134 and F167 conferring benzimidazole resistance. Only ASVs of hookworm as a percentage of the total are used and the sequencing read depth is depicted underneath. Samples that did not amplify hookworms are indicated by squares and samples with <2,000 sequence reads are depicted with stars. (C) Bar chart for Q134 and F167 isotype-1 β-tubulin of *A. caninum* where the proportion of an alternative residue coding for an amino acid that confers benzimidazole resistance are shown. Samples with no bar chart are those that did not amplify *A. caninum*, thus are no *A. caninum* ASVs.

### 3.2. Widespread and high frequency of A. caninum canonical F167Y (TTC>TAC) isotype-1 β-tubulin benzimidazole resistance mutation in Australia

For 81 Australian samples, the isotype-1 β-tubulin BZ167 amplicons included *A. caninum* isotype-1 β-tubulin with an average sequencing read count between 81 to 31,265 (Fig. 1B). Excluding six samples (TS6, TS7, TS39, TS55, TS63, TS90; sequence reads 81 to 1,699) with <2,000 sequences, the average was 11,092 sequences (*n*=75; 2,213 to 31,265). There were 72 samples with >2,000 isotype-1 β-tubulin BZ167 amplicon sequences belonging to *A. caninum* (average sequencing reads 11,271) (Fig. 1B).

Among the samples, 49% (35/72) contained the Q134H (CAA>CAT) resistant allele, with an average frequency of 10% (range 1% to 45%; 95% confidence interval (95% CI): 7-13%). Similarly, 67% (48/72) of the samples contained the F167Y (TTC>TAC) resistant allele, with an average frequency of 41% (range 1% to 100%; 95% CI: 33-51%) (Fig. 1C). There were 28% (20/72), 25% (18/72) and 14% (10/72) of samples with >40%, >50% and >75% F167Y (T<u>T</u>C>T<u>A</u>C) frequency, respectively (Fig. 1C). Moreover, 36% (26/72) of samples had both Q134H (CA<u>A</u>>CA<u>T</u>) and F167Y (T<u>T</u>C>T<u>A</u>C) benzimidazole-resistant alleles. The six samples with <2,000 sequences for *A. caninum* had only the susceptible (coding for Q134, F167) alleles, except TS90 with 100% of F167Y (T<u>T</u>C>T<u>A</u>C) mutation. Sequence analysis of isotype-1 β-tubulin sequence (amino acids E198 and F200; BZ200 amplicons) revealed no evidence of benzimidazole resistance signatures.

Besides *A. caninum* isotype-1 β-tubulin sequences, there were four *U. stenocephala* and one (1/4) of the samples TS24 included F167Y (T<u>T</u>C>T<u>A</u>C) in the isotype-1 β-tubulin mutation in 51% of the sequence reads. The remaining isotype-1 β-tubulin of *U. stenocephala* matched benzimidazole susceptible sequence residues, including across BZ200 amplicon (E198 and F200). The cat TS42 sample with *A. tubaeformae* and its isotype-1 β-tubulin included only the benzimidazole-susceptible sequences (amino acids Q134, F167, E198 and F200).

For the New Zealand samples only 50% (10/20) of DNA yielded C_t_<35 (20.2 to 33.6); nine samples yielded ITS-2 rDNA sequence data to consider species identification (TS76-TS84; average 18,488; 1,118 to 27,624 sequencing reads) (Fig. 1A). Four of the nine samples contained sequence data that belonged to non-hookworm sequence including ITS-2 rDNA sequencing matched to parasites of sheep and rabbits; three DNA samples contained only ASVs that matched spurious parasites (=‘pseudoparasite’) DNA sequences. Subtracting the ‘pseudoparasite’ sequences to retain hookworm sequence data revealed the presence of uniquely *U. stenocephala* sequences, five samples with 8,027 to 27,624 sequencing reads belonging to hookworms and one sample (TS76) with only 118 sequences. Sequence analysis of the isotype-1 β-tubulin (amino acids Q134, F167, E198 and F200) revealed no evidence of benzimidazole resistance signatures in New Zealand samples.

### 3.3. High frequency of Q134H (CAA>CAT) and F167Y (TTC>TAC) mutations in A. caninum in greyhounds

We then focused on 55 dog samples from two diagnostic laboratories (Vetnostics, VPDS) for which we had *A. caninum* Q134H (CA<u>A</u>>CA<u>T</u>) and F167Y (T<u>T</u>C>T<u>A</u>C) mutation frequency in the isotype-1 β-tubulin. There were 53 dogs with a defined location to a postcode from New South Wales, Australia (one dog from New South Wales did not have a postcode and the other dog was from Queensland). Dogs came from 12 distinct New South Wales postcodes (Fig. 2). The *A. caninum* faecal egg count across all 55 dogs was 534 EPG (minimum 20, maximum 4,920)

**Fig. 2.**
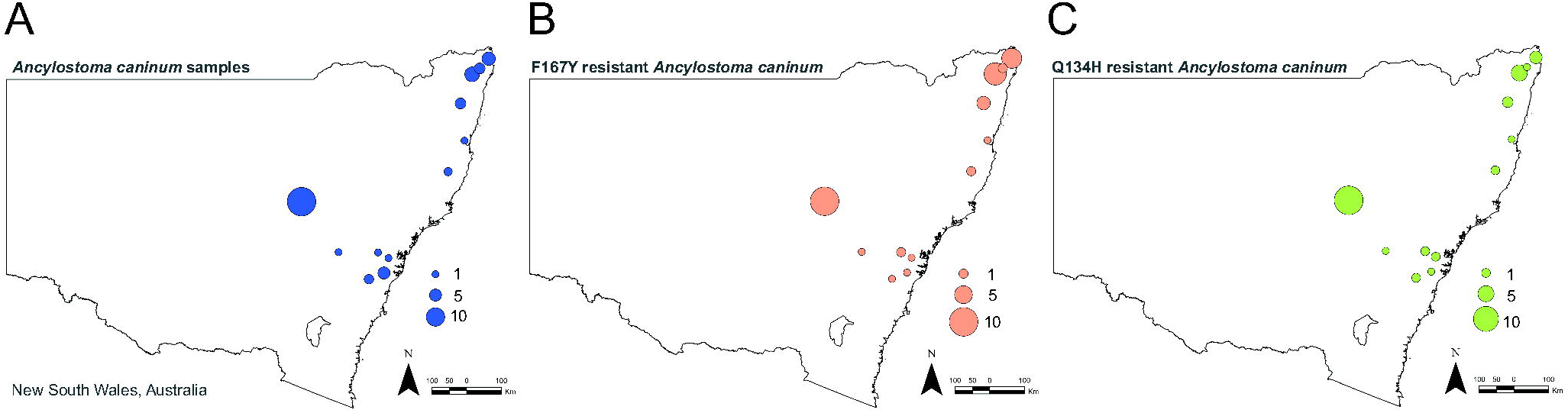
Distribution of *Ancylostoma caninum* and benzimidazole-resistant isotype-1 β-tubulin single nucleotide polymorphism. Proportional symbol maps of (A) the number of *A. caninum*-positive samples, and (B) number of Q134H (CA<u>A</u>>CA<u>T</u>) and (C) number of F167Y (T<u>T</u>C>T<u>A</u>C) isotype-1 β-tubulin of *A. caninum-*positive samples. Samples were summed per postcode in New South Wales, Australia. Samples are joined to postcodes of Australia shapefile (Geocentric Datum of Australia, GDA 1994) in ArcGIS^®^ Pro.

Greyhounds represented 80% (44/55) of the samples (42 from New South Wales, one from Queensland, one from an unknown state). The F167Y (T<u>T</u>C>T<u>A</u>C) allele was present in 30 (68%, 30/44) of them with 37% average frequency (mininum 2%, maximum 100%). The Q134H (CA<u>A</u>>CA<u>T</u>) allele was present in 28 (64%,28/44) of them with 10% average frequency (minimum 1%, maximum 45%).

There was no significant correlation between Q134H (CA<u>A</u>>CA<u>T</u>) or F167Y (T<u>T</u>C>T<u>A</u>C) and EPG (correlation coefficient=-0.169, *P*=0.217; correlation coefficient=0.03, *P*=0.82). Considering the association of presence or absence of either Q134H (CA<u>A</u>>CA<u>T</u>) or F167Y (T<u>T</u>C>T<u>A</u>C) and EPG, there was no significant association for either SNP (Kruskal-Walis Test: H=0.126, degrees of freedom (df)=1, *P*=0.72; H=0.56, df=1, P=0.45). We then considered the least stringent threshold of >40% for F167Y (T<u>T</u>C>T<u>A</u>C) mutation and recalculated the association, but the test remained not significant (Kruskal-Walis Test: H=0.02, df=1, *P*=0.89). There was only a single sample with >40% of the Q134H (CA<u>A</u>>CA<u>T</u>) mutation, therefore preventing calculation of these test statistics. Considering the ages of the dogs (for those available, *n*=41) there was no significant correlation between age and EPG, Q134H (CA<u>A</u>>CA<u>T</u>) or F167Y (T<u>T</u>C>T<u>A</u>C) (Spearman correlation, *P*>0.05).

The median resistance allele frequency in *A. caninum* samples from greyhounds for both the Q134H (CA<u>A</u>>CA<u>T</u>) and F167Y (T<u>T</u>C>T<u>A</u>C) mutation (0.035 and 0.090, respectively) was significantly higher than that for other dog breeds (median 0.000; Kruskal Wallis Test, *P*≤0.01) (Fig 2). There was no significant difference between EPGs for the two breed categories (Kruskal-Wallis Test, *P*=0.22).

### 3.4. Phenotypic evidence of benzimidazole resistance in A. caninum field sample with >75% F167Y (T<u>T</u>C>T<u>A</u>C) mutation

We tested 13 *A. caninum* samples (D1, D2, D3, D5, D6, D8, D9, D10, D12, D15, D16, D17, D18/20) with EHAs that had their frequency of Q134H (CA<u>A</u>>CA<u>T</u>) and F167Y (T<u>T</u>C>T<u>A</u>C) *A. caninum* SNP established (Fig. 1, Table 1). Dose response curves and IC_50_ formed two groups, those that had high IC_50_>0.15 µg/mL and those that had low IC_50_<0.15 µg/mL of thiabendazole (Fig. 3A). Three greyhounds (D1/D2: TS87/88, D3: TS89, D17: TS103) and the crossbreed dog (D6: TS92, D18/20: TS104/106) were in the high IC_50_ group and they all had >75% of F167Y (T<u>T</u>C>T<u>A</u>C) *A. caninum* mutation in the isotype-1 β-tubulin (Fig. 1, Table 1). The IC_50_ of the remaining seven greyhound samples ranged from 0.02 to 0.06 µg/mL of thiabendazole and the frequency of *A. caninum* F167Y (T<u>T</u>C>T<u>A</u>C) was <40% (Fig. 1, Table 1). For one dog (represented by two samples, D6 and D18/20, in Fig. 1), we established an in vitro EHA on eggs purified from the dog’s faeces on day 1 and day 9, showing an IC_50_ of 0.24 (0.19 to 0.30) µg/mL and 0.17 (95%CI 0.12 to 0.24) µg/mL of thiabendazole, respectively. There was no significant difference between the response curves for these two samples, demonstrated by a single response curve fitting both sample responses to thiabendazole (*P*=0.19; F=1.735, DFn=2, DFd=45). The benzimidazole susceptible control IC_50_ was 0.11 (95%CI 0.10 to 0.12) µg/mL (Fig. 3), and the *A. caninum* DNA was devoid of canonical F167Y (T<u>T</u>C>T<u>A</u>C) and Q134H (CA<u>A</u>>CA<u>T</u>) isotype-1 β-tubulin mutations which confer benzimidazole resistance; the BZ167 and BZ200 amplicons coded for amino acids Q134, F167, E198 and F200. There was significant correlation of the IC_50_ for thiabendazole in EHAs and F167Y (T<u>T</u>C>T<u>A</u>C) mutation in the isotype-1 β-tubulin (Spearman’s r=0.83 (95% CI=0.52 to 0.95, *P* < 0.001) (Fig. 3B).

**Fig. 3.**
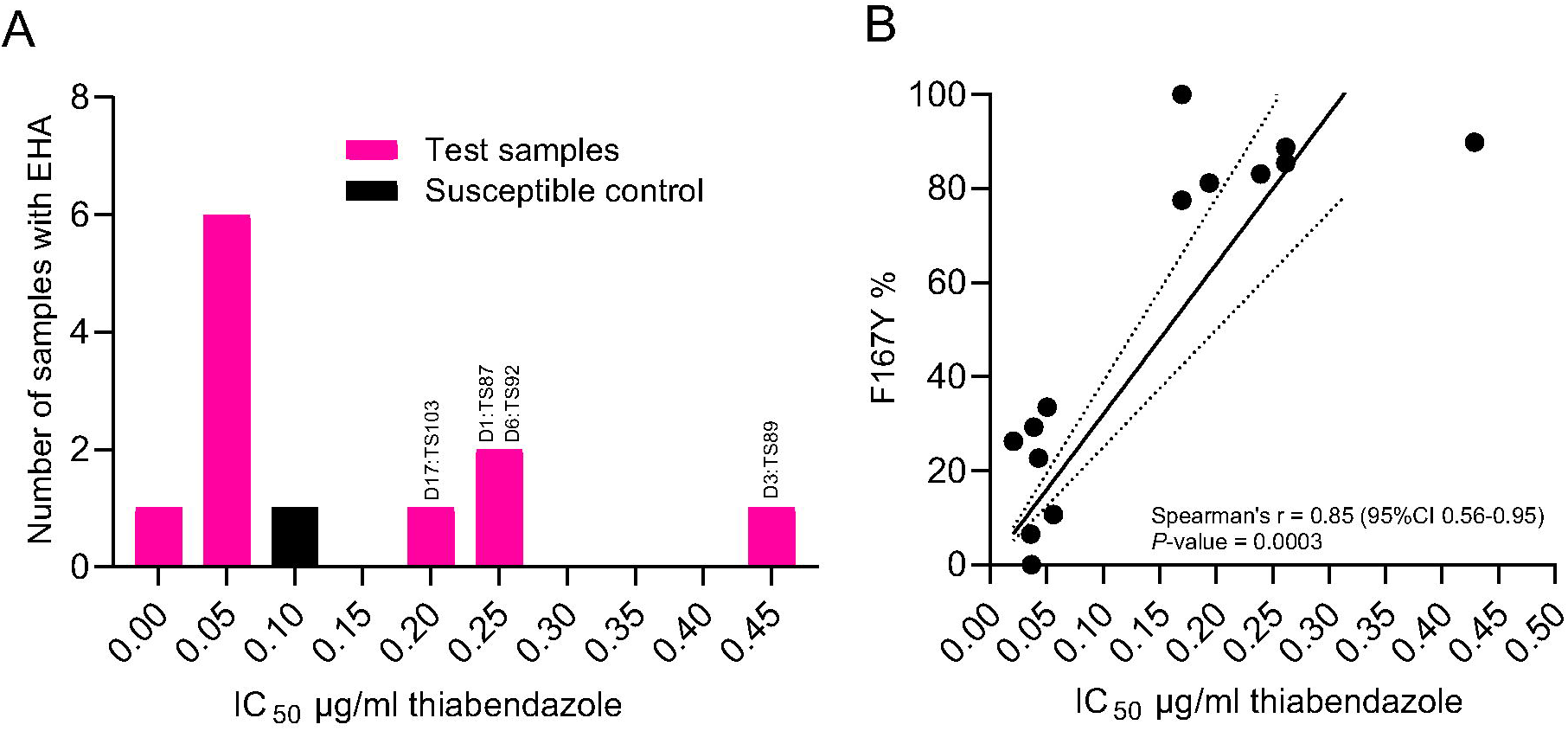
*Ancylostoma caninum* 50% inhibitory concentration (IC_50_) for thiabendazole in egg hatch assays. (A) Correlation IC_50_ values for *A. caninum* samples for thiabendazole agar-based egg hatch assay (EHA) from Queensland, Australia with the percentage of F167Y (T<u>T</u>C>T<u>A</u>C) isotype-1 β-tubulin of *A. caninum* present in the sample. Simple linear regression is depicted as a solid line with 95% confidence interval (95% CI) depicted as a dotted line. (B) Histogram of IC_50_ values for *A. caninum* samples from Queensland, Australia. A susceptible *A. caninum* sample, that was confirmed to have 100% faecal egg count (FEC) reduction, is colour coded (black). The samples with >0.15 µg/mL thiabendazole IC_50_ values have their identifiers above the bars.

**Table 1.** Summary of *Ancylostoma caninum* egg hatch assay with frequency of Q134H and F167Y polymorphisms in isotype-1 β-tubulin.

| Sample ID | Breed | EPG | IC<sub>50</sub> (µg/mL)<sup>a</sup> | IC<sub>50</sub> 95% CI (µg/mL)<sup>a</sup> | Q134H<sup>c</sup> | F167Y<sup>c</sup> |
| --- | --- | --- | --- | --- | --- | --- |
| High IC<sub>50</sub>>0.15 µg/mL of thiabendazole |  |  |  |  |  |  |
| D3:TS89 | Greyhound | 1,275 | 0.429 | 0.344 to 0.534 | 0% | 89.91% |
| D1:TS87<sup>d</sup> | Greyhound | 1,050 | 0.298 | 0.216 to 0.412 | 0% | 85.47% |
| D2:TS88<sup>d</sup> | Greyhound | 645 | 0.262 | 0.237 to 0.289 | 0% | 88.80% |
| D6:TS92<sup>e</sup> | Crossbreed | 540 | 0.239 | 0.201 to 0.285 | 0% | 83.11% |
| D18/D20:TS104/106<sup>e</sup> | Crossbreed | 225 | 0.170 | 0.121 to 0.239 | 0%/0% | 100%/77.54% |
| D17:TS103 | Greyhound | 5,535 | 0.194 | 0.150 to 0.250 | 2.55% | 81.23% |
| Low IC<sub>50</sub><0.15 µg/mL of thiabendazole |  |  |  |  |  |  |
| D5:TS91 | Greyhound | 14,910 | 0.056 | 0.048 to 0.066<sup>b</sup> | 0% | 10.69% |
| D9:TS95 | Greyhound | 930 | 0.050 | 0.043 to 0.059<sup>b</sup> | 18.27% | 33.51% |
| D10:TS96 | Greyhound | 4,080 | 0.043 | 0.040 to 0.046<sup>b</sup> | 0% | 22.67% |
| D8:TS94 | Greyhound | 390 | 0.039 | 0.031 to 0.048<sup>b</sup> | 0% | 29.26% |
| D12:TS98 | Greyhound | 3,930 | 0.037 | 0.035 to 0.039<sup>b</sup> | 0% | 0% |
| D15:TS101 | Greyhound | 720 | 0.036 | 0.031 to 0.041<sup>b</sup> | 0% | 6.56% |
| D16:TS102 | Greyhound | 885 | 0.021 | 0.013 to 0.033<sup>b</sup> | 7.43% | 22.53% |

<sup>a</sup> Thiabendazole concentration in µg/mL.
<sup>b</sup> Drug-susceptible *A. caninum* - 0.046 and 0.048 µg/mL (Jimenez Castro et al., 2019) and 0.045 µg/mL (Diawara et al., 2013).
<sup>c</sup> Percentage of canonical benzimidazole resistance codons on the isotype-1 β-tubulin.
<sup>d,e</sup> Same dog.
EPG, *A. caninum* eggs per gram of faeces; IC<sub>50</sub>, 50% inhibition concentration; 95% CI, 95% confidence interval.

In vivo evidence based on repeated FECs for one dog (D6 and D18/20 in Fig. 1) suggested decreased (<95%) ability of febantel/pyrantel to reduce egg shedding post-treatment. This crossbreed dog (D6, D18/20) initially shed 540 EPG of *A. caninum* hookworms on day 1 (day of treatment). The hookworm isolate carried 83% of F167Y (T<u>T</u>C>T<u>A</u>C) mutation in the isotype-1 β-tubulin (D6: TS092; Fig. 1), and was treated with ‘Drontal Allwormer’, but the faecal egg reduction only reached 88.8% (60 EPG) and 72.2% (195 EPG) on day 10 and day 14 post-treatment, respectively. The same dog was re-treated with macrocyclic lactone (‘Credelio Plus’) on day 20 (90 EPG) post original treatment, which resulted in 100% *A. caninum* egg reduction in faeces on day 8 post second treatment (0 EPG).

## 4. Discussion

This survey of dogs from south-eastern Australia infected with hookworms demonstrated a high frequency of *A. caninum* with canonical mutations F167Y and Q134H in the isotype-1 β-tubulin that are known to be associated with benzimidazole resistance. Both F167Y and Q134H mutations have been conclusively confirmed to lead to benzimidazole resistance using *Caenorhabditis elegans* as a surrogate (Kitchen et al., 2019; Venkatesan et al., 2023). The extent of both mutations, 67% for F167Y and 49% for Q134H, in *A. caninum* tested in this study exceeds even the recent USA report by Venkatesan et al. (2023) that utilised the same methodology based on NGS amplicon metabarcoding. Our samples from New South Wales correspond to hookworm-positive samples tested by a commercial diagnostic laboratory (*n*=49) over a 13-month period. Our sampling effort for New South Wales represents approximately 1,700 dog samples, assuming a conservative *A. caninum* prevalence of 2.9% in temperate and subtropical regions of Australia (Massetti et al., 2022).

Using our samples from New South Wales, we show that the isotype-1 β-tubulin F167Y mutation is present in canine hookworms across the state and predominantly in greyhounds that provided most of the samples. In North America, the isotype-1 β-tubulin F167Y mutation was present in all geographic areas with detected hookworms, including Canada (Leutenegger et al., 2024). Initially the F167Y polymorphism was recognised in *A. caninum* infecting greyhounds; currently the highest isotype-1 β-tubulin F167Y mutation prevalence was shown to be in the hookworms infecting poodles (29%), followed by Bernese mountain dogs (25%), cocker spaniels (23%) and greyhounds (22%) in North America (Jimenez Castro et al., 2019; Kitchen et al., 2019; Leutenegger et al., 2024). Using faecal samples from 17,671,724 individual dogs in North America, greyhounds were found to have a significantly higher risk of testing positive for hookworms on faecal flotation (odds ratio = 15.3) (Burton et al., 2024). Additionally, the interval to test negative was 71– 72 days for presumably anthelminthic-treated greyhounds, compared with just 1–2 days for other breeds (Burton et al., 2024). In this study from Australia, in addition to greyhounds, one crossbreed dog from Queensland was infected with *A. caninum* and had 83% of the isotype-1 β-tubulin F167Y mutation, together with a reduced in vitro IC_50_ for thiabendazole.

The EHA tests for in vitro susceptibility to benzimidazoles, thus phenotypes including *A. caninum* resistance to benzimidazole (Kotze et al., 2005, 2009; Diawara et al., 2013). Importantly, we demonstrated correlation of the EHA and the isotype-1 β-tubulin F167Y mutation, and those samples with elevated EHAs (IC_50_) all showed frequencies of F167Y mutation >75%. Known in vitro resistant *A. cainum* with an IC_50_of 8.76 μM (=1.76 µg/mL for KGR isolate) and 40.08 μM (=8.07 µg/mL for BCR isolate) for thiabendazole had 100% and 76% of the F167Y mutation (McKean et al., 2024). Our resistant *A. caninum* had an IC_50_ from 0.17 to 0.43 µg/mL of thiabendazole and are closer to the range (0.55 to 0.67 µg/mL) detected for resistant *A. caninum* by Jimenez Castro et al. (2019). The in vitro resistance was further confirmed by FEC reduction in one of the Australian dogs in vivo. In a study by Jimenez Castro et al. (2021), no significant correlation was found between the F167Y (TTC>TAC) mutation frequency and the IC_50_ for thiabendazole in a small study involving 15 *A. caninum* samples. However, three samples with an in vitro resistance phenotype exhibited very low (<6%) F167Y (TTC>TAC) mutation frequency (Jimenez Castro et al., 2021). This discrepancy was later explained by the discovery of a new mutation in isotype-1 β-tubulin, Q134H (CAA>CAT), which was present at a high frequency (≥90%) in all three in vitro resistant *A. caninum* samples studied by Venkatesan et al. (2023).

The detection of the isotype-1 β-tubulin F167Y mutation in *U. stenocephala*, the Northern hookworm that commonly infects dogs in cooler climates, is unexpected, as drug resistance has not previously been reported for *U. stenocephala* (Leutenegger et al., 2023; Venkatesan et al., 2023; Stocker et al., 2024). In our samples from Australia, we detected *U. stenocephala* F167Y mutation in one out of four dog samples. More frequently in Australia, *U. stenocephala* is a parasite of red foxes which likely serves as the reservoir (Ryan, 1976). In a study of 930 foxes from New South Wales, the most prevalent nematodes were *Toxocara canis* (35.2%) and *U. stenocephala* (30.6%), while *A. caninum* was detected in only 70 (7.5%) foxes (Ryan, 1976). Although *U. stenocephala* is often considered relatively non-pathogenic, it has been acknowledged as a cause of subacute enteritis and hypoproteinaemia in dogs (Gibbs, 1958; Miller, 1968). The significance of the F167Y mutation in *U. stenocephala* in conferring benzimidazole resistance in the presence of a large *refugium* in red foxes will require further confirmation from in vitro and in vivo drug response studies.

The absence of resistant SNPs that would code for E198 and F200 in the isotype-1 β-tubulin of *A. caninum* in Australia agrees with reports from the USA (Venkatesan et al., 2023). The F200Y SNP has been documented once in *A. caninum* dog samples from Brazil, but the in vitro or in vivo phenotype was not measured (Furtado et al., 2014). The F200Y was detected in human *N. americanus* from Haiti with frequencies of 20-90%, and in an in vitro EHA assay these hookworm isolates were less sensitive to thiabendazole (IC_50_ = 0.12 to 0.26 µg/µL) (Diawara et al., 2013). Molecular dynamic simulations using isotype-1 β-tubulin showed that *A. caninum* and *A. ceylanicum* both shared the same drug binding profile, while *A. duodenale* and *N. americanus* both had different binding profiles with no overlap except for the E198 interaction that is shared with all nematodes (Jones et al., 2022).

With growing evidence of anthelmintic resistance in *A. caninum*, the question arises whether the fitness and ability of resistant strains to spread across the dog population differ from non-resistant strains. Recently, a strain of triple-resistant *A. caninum* (BCR) was shown to exhibit reduced ‘larval activation’, an initial step in the transformation of infective non-feeding larvae into migrating and feeding larvae in a permissive host (McKean et al., 2024). Larval activation in vitro is initiated when infective L3s (iL3s) of *A. caninum* are incubated with a host-mimicking signal such as dog serum to resume feeding over a period of 24 h (Hawdon and Schad, 1990; Hotez et al., 1993). The consequences of the 40-50% reduced initial activation of the *A. caninum* triple-resistant strain (BCR) reported by McKean et al. (2024) are not yet known but suggest a possible reduced ability to establish infection. However, we did not find any association between the SNPs and FEC in the dogs we studied. Our study does not permit the examination of relationships between FECs and other variables (e.g., worm fitness) because the infection histories (such as exposure to larvae and drug treatment history) of the dogs are completely unknown. Our samples were neither tested for larval activation nor for the presence of pyrantel and macrocyclic lactone resistance, so reduced ‘larval activation’ may either be restricted to the triple-resistant strains, or it may not lead to lower infection rates and changes in egg output.

The molecular tools employed in the present study enable agile moderate to high throughput tracking of anthelmintic resistance mutations. This offers advantages in terms of speed and cost compared with phenotypic in vivo and in vitro measurements of sensitivity to anthelmintic drugs. We identified the hookworm species and simultaneously discovered widely distributed isotype-1 β-tubulin SNPs across eastern Australia. Hookworms, whipworm and roundworms in dogs and cats are traditionally managed by sustained routine administration of anthelminthics. The recent discovery of MADR in canine hookworm (*A. caninum*) has raised concerns within the veterinary profession (von Samson-Himmelstjerna et al., 2021; Marsh and Lakritz, 2023). Adopting a risk assessment-based approach to limit unnecessary use of parasiticides has the potential to mitigate the risk of resistance selection and its spread, and should be an important consideration in responsible parasite control (Selzer and Epe, 2021; Bagster and Elsheikha, 2022).

## Acknowledgements

Funding was provided by the Margo Roslyn Flood Bequest (Sydney School of Veterinary Science, The University of Sydney, Australia), and by The Canine Research Foundation and Dogs Victoria (School of Veterinary Science, University of Queensland, Australia).

